# *In vitro* longitudinal lumbar spinal cord preparations to study sensory and recurrent motor microcircuits of juvenile mice

**DOI:** 10.1101/2022.04.25.489385

**Authors:** Mustafa Görkem Özyurt, Julia Ojeda-Alonso, Marco Beato, Filipe Nascimento

## Abstract

*In vitro* spinal cord preparations have been extensively used to study microcircuits involved in the control of movement. By allowing precise control of experimental conditions coupled with state-of-the-art genetics, imaging and electrophysiological techniques, isolated spinal cords from mice have been an essential tool in detailing the identity, connectivity and function of spinal networks. The majority of the research has arisen from *in vitro* spinal cords of neonatal mice, which are still undergoing important postnatal maturation. Studies from adults have been attempted in transverse slices, however, these have been quite challenging due to the poor motoneuron accessibility and viability, as well as to the extensive damage to the motoneuron dendritic trees. In this work, we describe two types of coronal spinal cord preparations with either the ventral or the dorsal horn ablated, obtained from mice of different postnatal ages, spanning from pre-weaned to one month old. These semi-intact preparations allow recordings of sensory-afferent and motor-efferent responses from lumbar motoneurons using whole cell patch-clamp electrophysiology. We provide details of the slicing procedure and discuss the feasibility of whole-cell recordings. The *in vitro* dorsal and ventral horn-ablated spinal cord preparations described here are an useful tool to study spinal motor circuits in young mice that have reached the adult stages of locomotor development.

**New & Noteworthy:** In the past 20 years, most of the research into the mammalian spinal circuitry has been limited to *in vitro* preparations from embryonic and neonatal mice. We describe two *in vitro* longitudinal lumbar spinal cord preparations from juvenile mice, that allow the study of motoneuron properties and respective afferent or efferent spinal circuits through whole-cell patch-clamp. These preparations will be useful to those interested in the study of microcircuits at mature stages of motor development.

## Introduction

Initial investigations into the physiology of spinal circuits began more than 100 years ago with ground-breaking work on spinal reflexes. These earlier studies relied on recordings of muscle activity obtained from primates, cats, rabbits and other species, to elegantly illustrate the complexity of rhythmic patterns involved in movement and their dependence on sensory stimuli (Sherrington, 1906). Decades later, the introduction of sharp electrodes allowed to perform the first intracellular recordings from cat motoneurons *in vivo*, providing valuable information on the physiology of motoneurons and a first description of the organization of the circuits underlying spinal reflexes (Eccles, 1957; see Brownstone, 2006 and Hultborn, 2006). The cat *in vivo* preparation proved to be an invaluable tool in the study of spinal afferent and efferent systems, but is not amenable to pharmacological manipulations, nor it allows visualization of tissue and targeting of specific cells, other than those who could be functionally identified, like Ia inhibitory interneurons and Renshaw cells (Hultborn & Udo, 1972; Renshaw, 1946), that however proved to be very difficult to target systematically, due to their smaller size, compared with motoneurons. *In vitro* spinal cord preparations were initially developed in neonatal rats (Otsuka & Konihsi, 1974), mice (Bagust & Kerkut, 1979) and golden hamsters (Bagust et al., 1982). The *in vitro* conditions made possible the application of drugs, as well as a precise control of different experimental settings such as concentration of ions, pH and temperature, and later, also allowed for simultaneous Ca^2+^ imaging of a large number of cells (O’Donovan et al., 1994; Wilson et al., 2007), which permitted the design of experiments that could not be conceived *in vivo*. With the development of whole cell patch-clamp electrophysiology (Hamill et al., 1981), neuroscientists were also able to use isolated spinal cord preparations, either *en bloc* or in slices, to study the physiology of motoneurons and their synaptic inputs (Takahashi, 1990). The availability of genetically modified mice, with cells labelled with fluorescent proteins according to their genetic or neurotransmitter signature, made the *in vitro* spinal cord preparations from mice the benchmark tool for the study of mammalian spinal circuits.

Indeed, over the past two decades, the development of new genetic tools has largely broadened our understanding of mammalian spinal cord circuitry (Arber, 2012; Goulding, 2009). The identification of neuronal populations based on the expression of specific transcription factors (Jessell, 2000), allowed to selectively target or manipulate spinal neurons, prompting ground-breaking work into the understanding of spinal networks controlling movement. All these achievements and improvements are somewhat diminished by the ‘elephant in the room’ among spinal cord electrophysiologists: *in vitro* recordings from the spinal cord were mostly performed in embryonic and neonatal preparations, when motoneurons and motor circuits are only partially developed (Clarac et al., 2004; Sharples & Miles, 2021; Smith & Brownstone, 2020; Whelan, 2003). *In vitro* spinal cord preparations, and especially recordings from motoneurons, from late postnatal mice, have been challenging: motoneurons are very sensitive to anoxia and the thick layer of white matter enveloping the spinal cord prevents sufficient exchange of oxygenated solution within the tissue, leading to motoneuron death. On the other hand, while slices (especially the most commonly used transverse slices) grants direct exposure of the grey matter to the perfusion solution, motoneurons are often damaged during the slicing procedure due to their extensive dendritic arborization and often are not viable in mature sliced tissue. Consequently, hindlimb-innervating motoneurons from the few attempted *in vitro* preparations from adult mice, generally have very low survival rate with the majority of the recordings being from smaller size cells that tend to resist anoxia and mechanical stress (Hadzipasic et al., 2014; Mitra & Brownstone, 2012).To address this limitation, other spinal preparations from mice have been developed, such as *in vivo*-anaesthetized and decerebrated mouse spinal cord preparations which have been extremely useful in detailing motoneuronal function from adult mice with sharp electrode recordings (Manuel et al., 2009; Meehan et al., 2010; Meehan et al., 2017). However, the nature of these preparations heavily restricts the efficiency of genetic and imaging tools to study microcircuit function. Furthermore, electrophysiological recordings from motoneurons (to date) can only be performed using sharp electrodes, since the only access to motor nuclei is through the dorsal horn and blunt patch electrodes cannot penetrate that far. Despite technical difficulties (Jensen et al., 2021; Manuel, 2021; Manuel & Zytnicki, 2021), sharp recordings can be used to record membrane and firing properties, but they are not suitable for voltage clamp recordings, thus making it difficult to characterize synaptic conductances in an intact preparation. Recently, researchers have been able to successfully extend the age of *in vitro* motoneuron recordings from mice, to their juvenile period (P14-30), by tweaking the composition of the solutions used, relying on a very fast dissection and slicing at cold temperatures (Bhumbra & Beato, 2018; Sharples & Miles, 2021; Smith & Brownstone, 2020). These studies have been done in transverse spinal cord slices, an useful preparation to study motoneuron physiology, but that largely ablates motoneuron dendritic arborization and pre-motor networks (Mousa & Elbasiouny, 2021). To better understand spinal microcircuit function in juvenile mice, studies from more intact neuronal preparations that would preserve motoneurons *in vitro* while keeping pre-motor networks intact and accessible, are needed.

In this work, we performed whole-cell patch clamp recordings from motoneurons from two different longitudinal mouse lumbar spinal cord preparations, to show that is it possible to study *in vitro* pre-motor microcircuit physiology in mice as old as P36. We used: 1) a preparation in which we removed part of the ventral horn thus exposing dorsolateral L3-L5 motoneurons, allowing the study of sensory-related circuits through dorsal root stimulation; and 2) a longitudinal spinal cord section with part of the dorsal horn removed, allowing to record inputs from recurrent motor efferent pathways to motoneurons following ventral root stimulation. Since the mice used here (P14-36) are past the weight-bearing stages and display motor behaviours associated with adulthood (Altman & Sudarshan, 1975), with motoneurons and their sensory and motor pathways fully developed (Alvarez et al., 2013; Sharples & Miles, 2021; Siembab et al., 2010; Smith & Brownstone, 2020), the longitudinal *in vitro* spinal cord preparations we describe here will provide a useful tool to anyone interested in studying mammalian spinal microcircuits at more mature stages of development.

## Methods

### Animals

All the experiments described were performed in conformity with the UK Animals (Scientific Procedures) Act 1986 and were approved by University College London ethical review committees and abided to UK Home Office regulations. Experiments were performed on both male and female mice on a C57BL/6J background. In this study we used mice of different ages, from postnatal P14 to late juvenile (P36) periods. All mice were weaned by P21 and separated into same sex cages.

### Intramuscular injections

In a subset of experiments we labelled motoneurons innervating the dorsiflexor tibialis anterior and/or plantar flexor gastrocnemius muscles by intramuscular injections of Cholera Toxin subunit B (CTB) conjugated to a fluorescent dye (Fig. 1A). Injections were performed 2-5 days prior to electrophysiological recordings. Animals were anaesthetized with isofluorane and regularly checked during the injection procedure for absence of tail and paw pinch reflexes. The hindlimb was immobilised and a small incision made through the skin and deep fascia to expose the muscle. A thin glass needle (30-50 μm diameter) loaded onto a Hamilton syringe, was used to slowly inject 1μL of CTB-Alexa-Fluor-488 or CTB-Alexa-555 (0.2% w/v in 1× phosphate buffer saline) into the belly of the tibialis anterior or lateral head of gastrocnemius muscles, respectively. The skin was then sutured and the animal allowed to recover from anaesthesia in a heated (30°C) incubation chamber.

**Figure 1.**
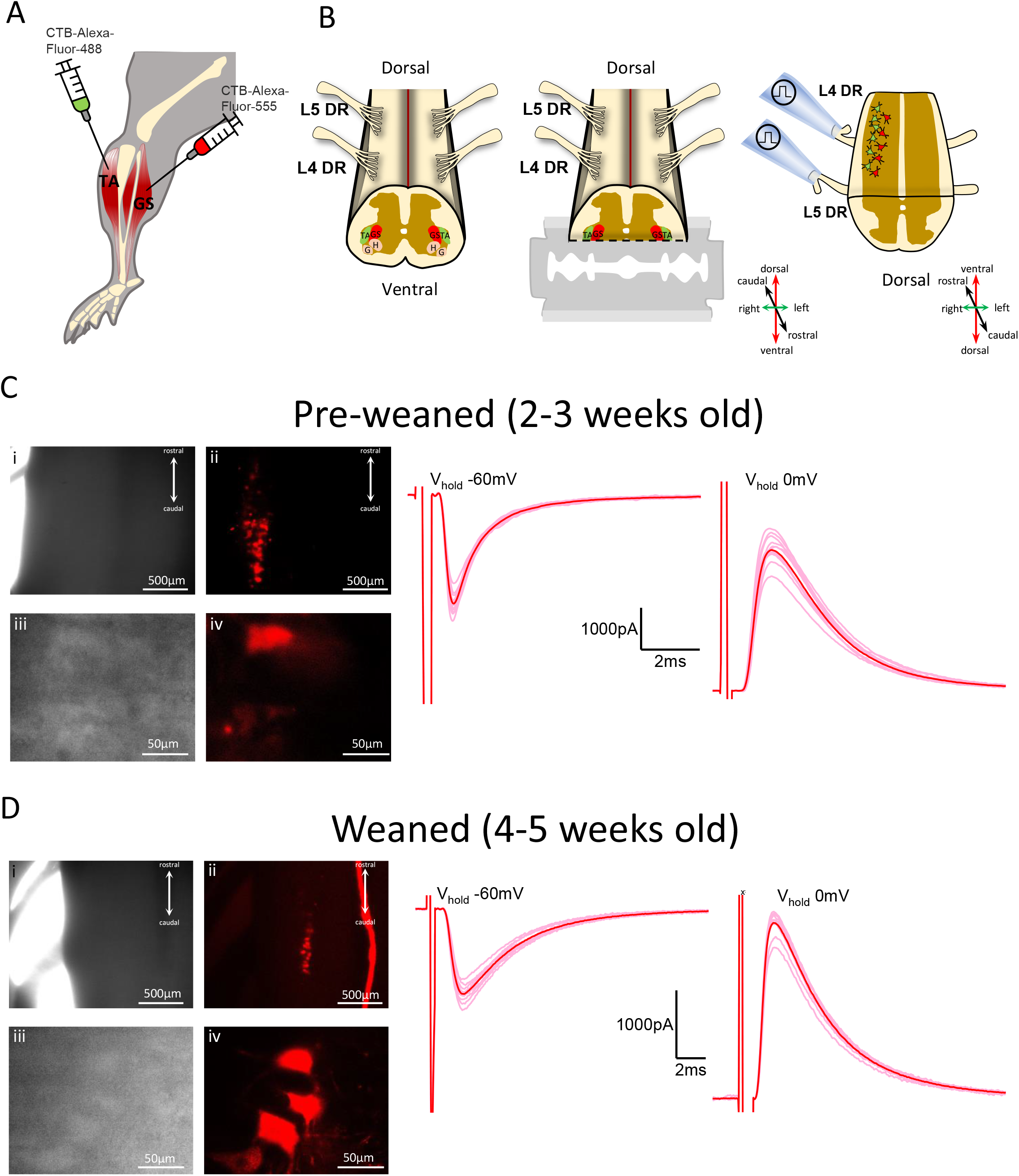
*In vitro* ventral horn-ablated spinal cord preparation allows electrophysiological access to lumbar motoneurons and their sensory-afferent circuits. **(a)** Schematic of the intramuscular injection performed to label the dorsiflexor tibialis anterior (TA) and/or gastrocnemius (GS) hindlimb muscles, and **(b)** representation of the isolated lumbar spinal cord with dorsal side facing up (left) and the coronal slicing performed to remove part of the ventral horn (middle), thus exposing the dorsolateral motor column and keeping dorsal roots (DR) intact for stimulation (right). H – hamstring, G – gluteus. DIC (grayscale) and confocal fluorescence (colored) images of the spinal cord preparation (top x4, bottom x40 magnification) from **(c)** pre-weaned 2-3 weeks old (P19) and **(e)** weaned 4-5 weeks old (P30) mice with gastrocnemius (in red) motor nuclei labelled, next to examples of excitatory (V_hold_ −60mV) and inhibitory (V_hold_ 0mV) postsynaptic currents recorded from animals of matching age.

### *In vitro* longitudinal spinal cord preparations

Mice were anesthetized with a mixture of ketamine/xylazine (100mg/kg and 10mg/kg, respectively) and decapitated. The spinal column was removed and pinned ventral side up in a chamber filled with recording artificial cerebrospinal fluid (aCSF) at ~2-4°C and composed of (in mM): 113 NaCl, 3 KCl, 25 NaHCO_3_, 1 NaH_2_PO_4_, 2 CaCl_2_, 2 MgCl_2_, and 11 D-glucose continuously bubbled with 95% O_2_ and 5% CO_2_. After laminectomy, the lumbosacral spinal cord was glued longitudinally to an agar block (7%) prepared with distilled water and 0.1% methyl blue, to increase the contrast and facilitate identification of landmarks in the spinal cord. Dorsal or ventral L3-L5 dorsal were left intact: for ventral horn-ablated preparations, the dorsal side of the cord was facing up (Fig. 1B, left), while for dorsal horn-ablated preparations, the ventral side was facing up (Fig. 2B, left). An oblique cut (~45° degrees) performed with surgical scissors above the L3 region, allowed visualization of the central canal under a dissection microscope mounted on top of the vibratome. The cord and agar were attached to a metallic specimen holder that was inserted into a vibratome chamber (HM 650V, Microm, ThermoFisher Scientific, UK) filled with ice-cold aCSF (~2°C) comprising (in mM): 130 K-gluconate, 15 KCl, 0.05 EGTA, 20 HEPES, 25 D-glucose, 3 Na-kynurenic acid, 2 Na-pyruvate, 3 Myo-inositol, 1 Na-L-ascorbate, pH 7.4 with NaOH (Dugué et al., 2005). The edge of the vibratome razor blade was aligned with the mid-point between the lower end of the central canal and the start of the ventral commissure white matter (for ventral horn removal, Fig. 1B, middle) or with the top of the central canal (for dorsal horn removal, Fig. 2A, middle). The spinal cord was slowly sectioned coronally (0.02 mm/s), thus obtaining a ventral or dorsal horn-ablated spinal cord preparation containing L3-L5 segments and roots intact (Figs. 1B, right and 2A, right). The tissue was incubated in a chamber with recording aCSF at 37°C for 30-45min, and then maintained at room temperature (20-22°C), constantly gassed with a mixture of 95/5% O_2_/CO_2_. Data shown were sourced from experiments performed at room temperature (20-22°C), aside from data from three dorsal horn-ablated preparations and six ventral horn-ablated preparations from 2-3 weeks old mice, which were collected at physiological temperature (31°C). The spinal cord of juvenile mice (≥2 weeks) is extremely susceptible to anoxia and mechanical stress, and therefore in order to obtain viable tissue for electrophysiological recordings we relied on a fast laminectomy and slicing, with the vibratome slicing starting within 10 minutes after decapitation, similar to our method for obtaining viable transverse slices from mature spinal cords (Bhumbra & Beato, 2018).

**Figure 2.**
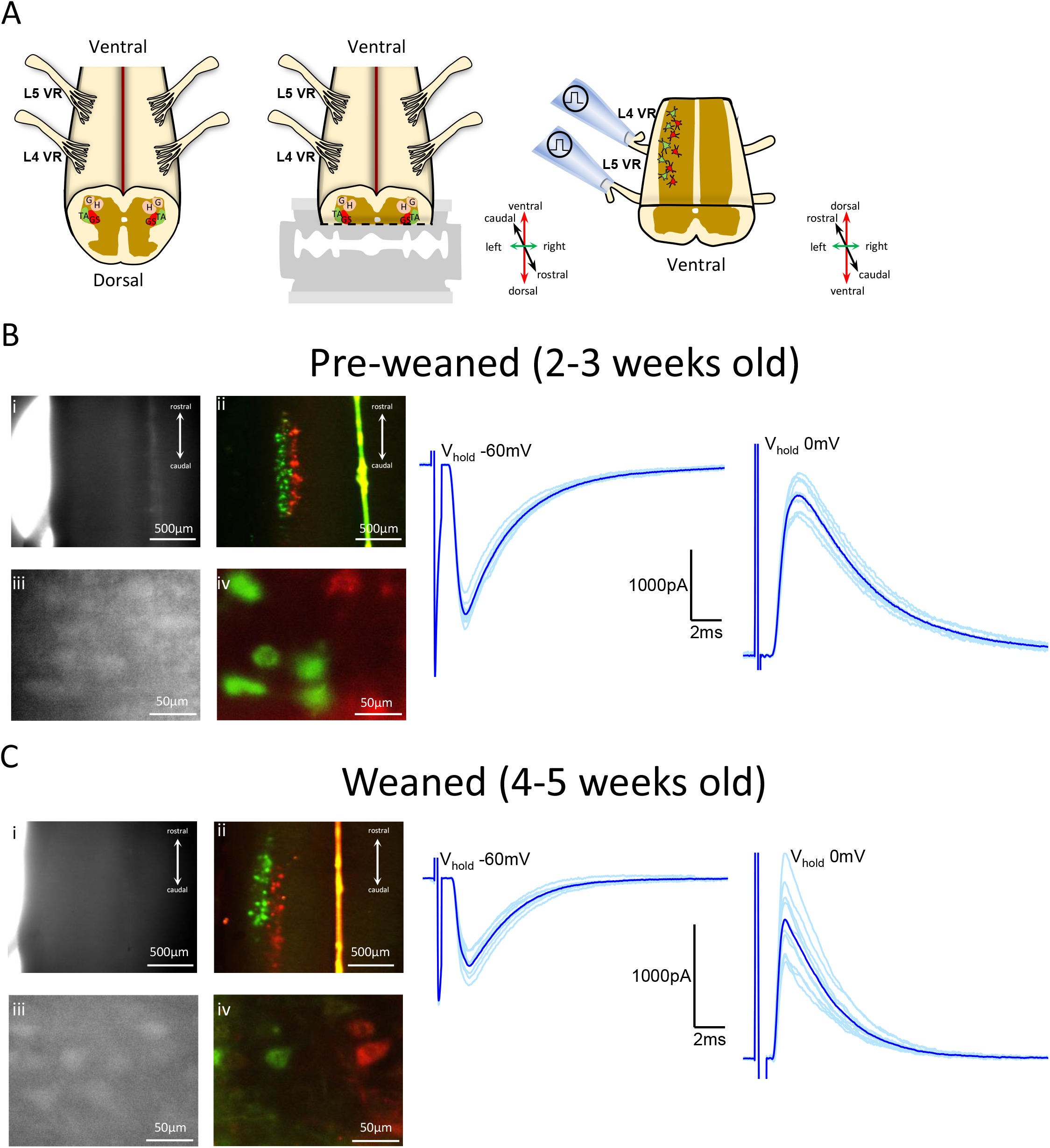
*In vitro* dorsal horn-ablated spinal cord preparation permits electrophysiological access to lumbar motoneurons and respective motor-efferent pathways. **(a)** Illustration of the isolated lumbar spinal cord with ventral side facing up (left) and the dorsal horn-ablation performed with the vibratome (middle), that allows to visualize the dorsolateral motor column and preserves ventral roots (VR, right). TA - tibialis anterior, GS – gastrocnemius. DIC (grayscale) and fluorescent (colored) imaging of spinal cord preparations (top x4, bottom x40 magnification) obtained from **(b)** pre-weaned 2-3 weeks old (P19) and **(d)** weaned 4-5 weeks old (P30) mice, with retrogradely labelled motoneurons (tibialis anterior in green and gastrocnemius in red), next to example traces of ventral root-evoked excitatory (V_hold_ −60mV) and inhibitory (V_hold_ 0mV) postsynaptic currents obtained from mice from respective age groups.

### *In vitro* imaging and electrophysiology

Motoneurons were distributed along the lateral rostro-caudal surface of both ventral and dorsal horn-ablated preparations and were visualized using an Eclipse E600FN Nikon microscope (Nikon, Japan) equipped with a double port that allowed simultaneous imaging of infrared differential interference contrast (DIC) images, through a digital camera (Nikon, DS-Qi1Mc), and fluorescence, detected through a laser scanning confocal unit (D-Eclipse C1, Nikon) equipped with two diode laser lines (λ=488 and 561 nm).

Whole-cell patch clamp recordings of motoneurons were performed using an Axopatch 200B amplifier (Molecular Devices, Sunnyvale, USA) and signals were filtered at 5kHz and acquired at 50kHz using a Digidata 1440A A/D board (Molecular Devices, Sunnyvale, USA) and Clampex 10 software (Molecular Devices, Sunnyvale, USA). Patch pipettes from borosilicate glass (GC150F, Harvard Apparatus, Cambridge, UK) were pulled with a Flaming-Brown puller (P1000, Sutter Instruments, CA, USA) to a resistance of ~1-4MΩ. Glass pipettes were filled with an intracellular solution containing (in mM): 125 Cs-gluconate, 4 NaCl, 0.5 CaCl_2_, 5 EGTA, 10 HEPES, 2 Mg-ATP, pH 7.3 with CsOH, and osmolarity of 290 to 310 mOsm. For six ventral horn-ablated preparations, data were acquired using a different intracellular solution composed of (in mM): 125 K-gluconate, 6 KCl, 10 HEPES, 0.1 EGTA, 2 Mg-ATP, with or without 3 QX-314-Br, pH 7.3 with KOH, and osmolarity of 290 to 310 mOsm.

Motoneuron capacitance and resistance were estimated from either the current change to a voltage step (5mV) in voltage clamp, or the voltage response to a brief (200ms) current step (50 to 100pA) in current clamp. Electrical stimulation was applied via a glass suction electrode, cut to size to fit tightly the diameter of the roots. Usually, both L4 and L5 ventral or dorsal roots were stimulated and the response to either root stimulation was recorded in the motoneurons. Stimulation was delivered by an isolated current stimulator (DS3, Digitimer, Welwyn Garden City, UK). The threshold intensity for evoking an initial response in the recorded motoneuron was determined and ventral roots were stimulated at 1.5-3x threshold, and dorsal roots at 3-5x threshold. Excitatory (EPSCs) and inhibitory postsynaptic currents (IPSCs) were recorded in voltage clamp mode at holding voltages of −60mV and 0mV, respectively, taking into account a correction for the junction potential (~15mV for both intracellular solutions), and their conductance was calculated from the size of the recorded current assuming a reversal of −60mV for inhibitory and 0mV for excitatory conductances. In a subset of recordings obtained from 2-3 weeks old mice, we estimated the conductance of recurrent inhibition in current clamp, in the presence of blockers of glutamatergic and GABAergic transmissions (D-2-amino-5-phosphonopentanoic acid - APV, 50μM; 1,2,3,4-tetrahydrobenzo(f)quinoxaline-7-sulphonamide - NBQX, 3μM; gabazine, 3μM), that is sufficient to fully abolish recurrent excitation without affecting recurrent inhibition received by motoneurons (Bhumbra & Beato, 2018). Synaptic conductance measurements were achieved by estimating the cell conductance from the linear fit of the voltage-current relationship in control and during the activation of the inhibitory synaptic conductances induced by 200 Hz VR stimulation (Suppl. Fig. 1A). Conductance of the recurrent inhibition estimated in either current or voltage clamp was similar (I_Clamp_ = 45±22nS, n = 66 root responses, V_Clamp_ = 52±42, n = 58 root responses; effect size mean difference = −7nS CI_95_ = [−19, 5]; Suppl. Fig. 1B).

### Statistical analysis

Raw traces were analyzed with Clampfit 10.7 (Molecular Devices, Sunnyvale, CA) and all statistical and data analyses were done using OriginPro 2021 (OriginLab Corporation, Northampton, MA, USA), Microsoft Excel version 2203 (Microsoft, Redmond, WA, USA), ImageJ/Fiji 1.53f51 (National Institutes of Health, Bethesda, Maryland, USA), MATLAB R2021a (Mathworks, Natick, MA) and R version 4.0.5 (R Core Team, Vienna, Austria). We used a linear mixed model (LMM) to examine the relationship between motoneuron capacitance, motoneuron conductance or root-evoked synaptic conductances and age of the preparation utilized. For this, we organized the data into two different age groups - preweaned (2-3 weeks old) and weaned (4-5 weeks old). The model takes into account the hierarchical dependence the data (Yu et al., 2022), which can be organized into 2 (animal and motoneuron) or 3 (animal, root and synaptic response) different levels. We used the *lmer* function contained in the package *Ime4* in R (Bates et al., 2015) to fit a LMM with a fixed-effect term for age group and a random intercept that varies by animal or root. The model was fitted as follows (Henderson et al., 1959; Yu et al., 2022):

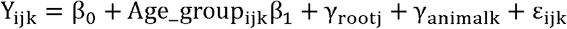

where Y_ijk_ corresponds to data for the i^th^ observation obtained from the j^th^ root (if applicable) from the k^th^ mouse; ß_0_ is the intercept; Age_group_ijk_ is the predictor variable for observation i grouped in j (if applicable) and k and β_1_ its coefficient; γ_rootj_ and γ_animalk_ are the random effects variables and ε_ijk_ is the observation error. The γ_rootj_ grouping variable was not used when fitting the LMM for capacitance because the data structure had one fewer level (measure of capacitance are unrelated to the stimulated root).

The data are shown as a scatter chart that displays each datapoint obtained for motoneuron capacitance, motoneuron conductance or synaptic conductances across age, juxtaposed to the LMM predicted difference between age groups shown as a dot and respective 95% confidence interval (CI) as a vertical bar, with both predictions aligned with respective groups by dotted horizontal lines (0 for pre-weaned and estimated difference for weaned mice, see Fig. 3A for example). CIs from estimated differences that do not include 0, would be considered as biologically relevant. In addition to the standard output of the LMM, we are also reporting partial eta squared (η_p_^2^) and its 90% CI on the top of the data graphs (Richardson, 2011; Steiger, 2004). Effect sizes provide a much better understanding of the magnitude of the differences found (Coe, 2002), and therefore the use of the η_p_^2^ estimate improves the comparability between groups. Regarding η_p_^2^, the effect size can be classified as small (0.01), medium (0.06) and large (0.14) (Cohen, 1988; Richardson, 2011). The η_p_^2^ can be obtained from the *t*-statistics from LMMs (*t*back method) (Correll et al., 2021). The values for η_p_^2^ were computed using the *t_to_eta2* function in the *effectsize* R package that uses the *t*-value and estimated degrees of freedom (df) to calculate η_p_^2^ as follows (Correll et al., 2021):

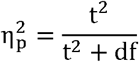

**Figure 3.**
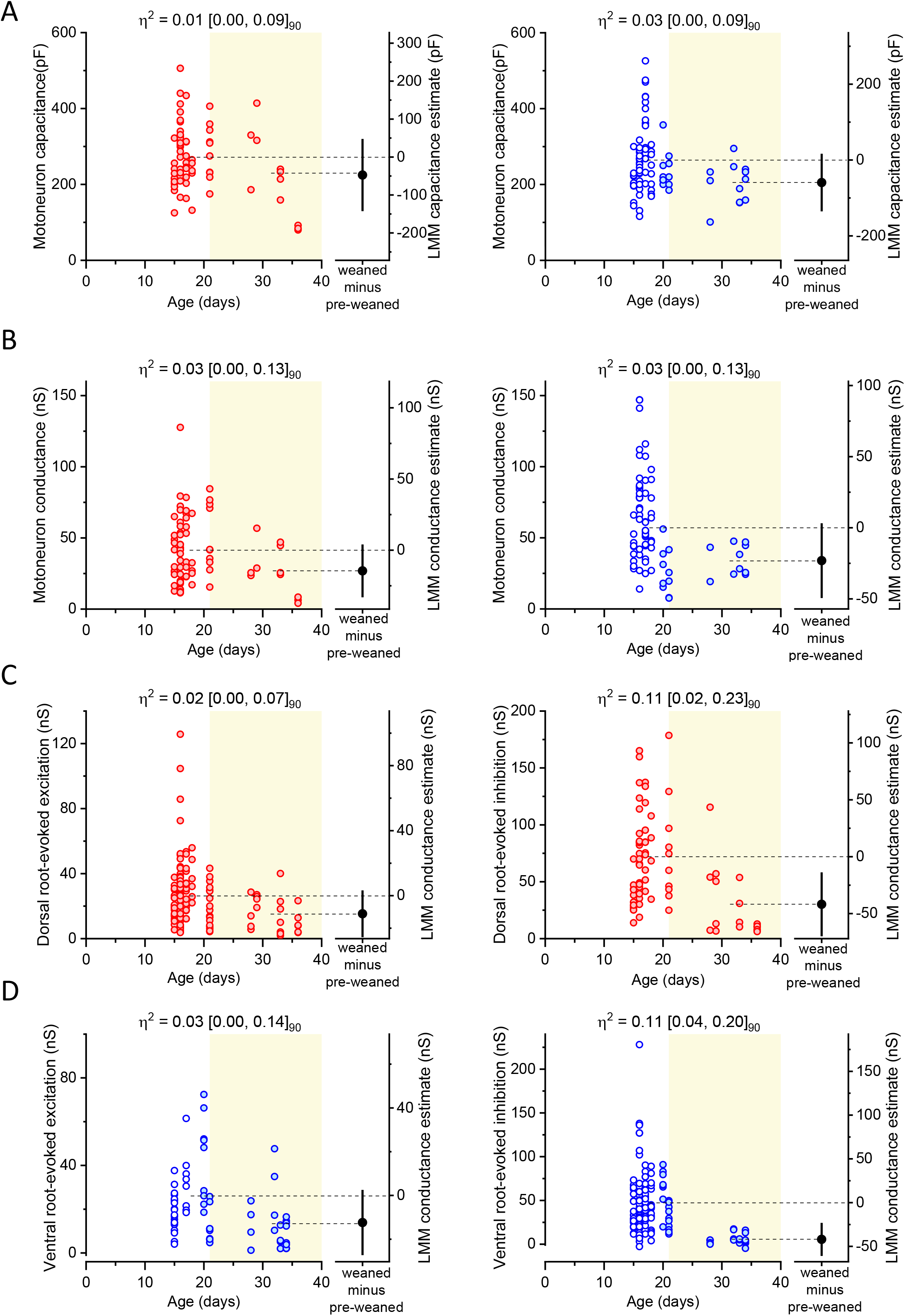
Motoneuron capacitance, motoneuron conductance and conductance of evoked synaptic currents obtained from *in vitro* ventral and dorsal-horn ablated spinal cords throughout age. Plots of individual data points plus LMM fixed-effect estimate difference between groups of **(a)** motoneuron capacitance and **(b)** conductance obtained from *in vitro* ventral- (red) or dorsal-ablated (blue) spinal cords across age, synaptic conductance for **(c)** dorsal root-evoked excitation (left) and inhibition (right) and synaptic conductance for **(d)** ventral root-evoked excitation (left) and inhibition (right) throughout age. Filled and non-filled dots represent experiments performed with Cs-gluconate or K-gluconate based intracellular solutions, respectively (see Methods). Light-yellow filled area was used to distinguish the datapoints corresponding to pre-weaned (P14-P21) and weaned (P22-P40) age groups. Partial eta squared (η_p_^2^) and respective 90% confidence interval are shown on top of each plot.

The output of the *t_to_eta2* function also estimates a 90% CI, appropriate for η_p_^2^ (Steiger, 2004). The variance (σ^2^) of the random effects given by the LMM is reported and used to calculate the intraclass correlation coefficient (ICC), which provides a partition of the sources of variability within our hierarchical structure (Yu et al., 2022).

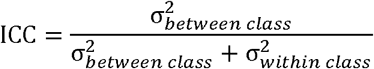

Comparisons between recurrent inhibition recorded in current and voltage clamp are depicted as estimation plots (Ho et al., 2019) with all individual points and respective boxplots shown on an upper panel, and a lower panel illustrating the effect sizes from a sampling distribution, 95% CI and mean difference obtained with bootstrapping (resampling done 10,000 times) performed in MATLAB. For bootstrapped mean comparisons, effect sizes that do not include 0, would be considered biologically significant (Ho et al., 2019). Standard descriptive stats describing the absolute mean ± standard deviation (SD) and number of observations (n) are also reported alongside parameters from LMM and effect sizes estimates.

## Results

To study sensory and motor-related spinal microcircuits from mice, we performed a longitudinal cut throughout the rostro-caudal axis of the L3-L5 mouse lumbar cord and removed either part of the ventral (Fig. 1B) or dorsal horn (Fig. 2A), in order to visualize and target motoneurons for electrophysiological recordings. Synaptic conductances from afferent or efferent pathways were then measured following dorsal or ventral root stimulation. The position of the cut gives electrophysiological access to the dorsal motor nuclei, located mostly in the L4 and L5 segments, innervating ankle flexor and extensor muscles. Nuclei identification was confirmed in a subset of experiments (ventral or dorsal horn-ablated), in which we labelled the lateral gastrocnemius and/or the tibialis anterior muscles through intramuscular injection of CTB conjugated with Alexa-Fluor-555 (red) and/or Alexa-Fluor-488 (green), respectively (Fig. 1A). We performed experiments on mice of different ages, from two weeks to five weeks old juvenile periods, encompassing a time-frame during which ongoing development and maturation impose different age-related technical constrains to obtaining viable *in vitro* preparations and recordings from motoneurons. Therefore, in order to appropriately convey the fundamental methodological aspects of *in vitro* recordings across age, we will be detailing the diverse practical aspects of experiments performed on dorsal and ventral horn-ablated preparations from: 1) 2-3 weeks old mice (pre-weaned), which have been recently utilized to obtain transverse slices and successfully record from lumbar motoneurons; and 2) 4-5 weeks old mice (weaned), which, along with 2-3 weeks old animals have reached advanced stages of locomotor development, but are beyond the age range used for viable whole-cell patch clamp motoneuron recordings in slices, and pose additional technical challenges to obtaining *in vitro* recordings from motoneurons.

### *In vitro* ventral horn-ablated lumbar spinal cord preparation

In order to record sensory-evoked inputs from motoneurons, we sectioned part of the ventral horn from the lumbar spinal cord of mice and kept dorsal roots intact for stimulation (Fig. 1B). As seen in figure 1C (left), the coronal slicing preferentially exposed dorsolateral motor nuclei such as those innervating the gastrocnemius (red). The exact position of the cut is critical: a cut that is too dorsal would reduce the number of preserved motoneurons if the blade goes through the nuclei, while a cut that is too ventral would not expose the motoneurons to the surface of the slice. Following dissection and slicing, the spinal cord was put under the microscope and the dorsal roots L4 and L5 were placed in a suction stimulating electrode.

We then tested the electrophysiological viability of motoneurons from longitudinal preparations. Motoneuron recordings from transverse slices have been already successfully performed in 2-3 weeks old pre-weaned juvenile mice (Bhumbra & Beato, 2018; Sharples & Miles, 2021; Smith & Brownstone, 2020). At this age myelination has progressed (Foran & Peterson, 1992) and the tissue becomes extremely dark, allowing poor detection of transmitted light under low magnification (Fig. 1C-i). However, the labelling of the gastrocnemius motor column was still visible in the low power confocal (Fig. 1C-ii). While some motoneurons on the immediate surface may not survive the cut, viable motoneurons were always found in the deeper layers. These could be barely identified within the infrared DIC image (Fig. 1C-iii), but were more apparent in the fluorescence channel (Fig. 1C-iv). Hence, some of these recordings were done partially blind since it became difficult to clearly visualize the contour of lumbar motoneurons at increased levels of depth. We stimulated the dorsal roots at an intensity of 1.5-3x the threshold for an initial evoked response, in order to preferentially activate the thickest fibers. This allowed us to record monosynaptic excitation received by motoneurons from Ia afferents (V_hold_ −60mV, Fig. 1C, left trace), as well as disynaptic inhibition (V_hold_ 0mV, Fig. 1C, right trace) known to derive from activation of Ia and Ib interneurons (Brink et al., 1983; Wang et al., 2021), proving that the afferent inputs to motoneurons and to Ia or Ib interneurons were anatomically intact and functional in this preparation. The large size of this preparation probably reduces tissue oxygenation. As a consequence, motoneurons were viable for only ~5-6 hours after the slicing recovery period.

While the preparations from mice younger than 3 weeks offer a good window for characterizing afferent circuits throughout postnatal development, we tested if we could extend the age of such experiments by recording from 4-5 weeks old weaned mice. At this age, the tissue is not transparent to infrared light (Fig. 1D-i) but labelling of the gastrocnemius motor column was visible under epi-fluorescence (Fig. 1D-ii). Under high magnification, the DIC image (Fig. 1D-iii) only partially revealed the shape of motoneurons, however targeted patch could be performed with the aid of the confocal image (Fig. 1D-iv, but images of similar quality can be obtained with a normal epifluorescence system), while in non-labelled preparations, motoneurons had to be patched semi-blind. We were able to record dorsal root-evoked Ia excitatory EPSCs and disynaptic inhibitory IPSCs (Fig. 1D, left and right traces respectively), albeit with a lower efficiency compared with that of younger preparations. The preparations from these older mice (P28-36) were not viable for long periods of time, and we usually could not get any motoneuron recording after more than 2h post-slicing recovery.

The *in vitro* ventral horn-ablated preparation thus allows access to motor nuclei innervating the lower limb, that can be identified and patched, with or without labelling, in spinal cords from pre-wean stages, to fully mature. Therefore, we could characterize not only the cellular properties of motoneurons, but, using dorsal root stimulation, we could also study sensory-related excitatory and inhibitory pathways up to an age in which sensory circuits are fully developed.

### *In vitro* dorsal horn-ablated lumbar spinal cord preparation

The ventral horn-ablated preparation described above gives full access to lower hindlimb motoneurons and preserves the integrity of afferent circuits originating in the dorsal horn. However it is not suitable for recording recurrent circuits, that are mostly located in the ventral horn. A dorsal horn ablated spinal cord preparation has already been described, and used in neonatal cords to record from V2a, Shox2, and Chx10 interneurons, as well as for fictive locomotion (Bhumbra & Beato, 2018; Dougherty & Kiehn, 2010; Li et al., 2019). Here we report that the dorsal horn-ablated preparation (Fig. 2A) can also be used to record from motoneurons innervating the lower limbs and to measure the synaptic inputs they receive from Renshaw cells and other motoneurons, evoked by antidromic stimulation of the ventral roots.

A preparation from a 2-3 weeks old animal is shown in figure 2B. Similarly to ventral horn-ablated preparations, dorsal horn-ablated spinal cords are dark and the infrared light has poor penetration (Fig. 2B-i). Even though this preparation is thinner than the ventral horn ablated one, the intensity of transmitted light is reduced because of the thickness of the white matter layer in the ventral horn and of the higher degree of myelination of ventral tissue. Also in preparations from these mice, the labelled motor columns (tibialis anterior in green and gastrocnemius in red) are clearly visible under low power confocal illumination (Fig. 2B-ii). Under higher magnification (Fig. 2B-iii) motoneurons were sufficiently visible in the infrared-DIC channel and those belonging to different nuclei could be identified in the fluorescence image (Fig. 2B-iv). Patch clamp recordings from identified motoneurons were visually guided and stimulation of the ventral root evoked an excitatory (Fig. 2B, left trace) and inhibitory (Fig. 2B, right trace) current, at their respective reversal potential. Similar to the ventral horn-ablated preparations from the same age range, the viability of pre-weaned P14-21 dorsal horn-ablated preparations extends to ~5-6 hours post-slicing recovery.

Finally, we also tested the feasibility of recording from motoneurons and measuring inputs from their recurrent circuits in *in vitro* preparations obtained from older, 4-5 weeks old, mice. Also in this case there was poor light penetration in the tissue (Fig. 2C-i), but gastrocnemius (red) and tibialis anterior (green) motor columns were clearly visible in the low power confocal image (Fig. 2C-ii). Under higher magnification the shape of motoneurons could still be detected in the infrared DIC image (Fig. 2C-iii), with clear labelling in the fluorescence channels (Fig. 2C-iv). In these animals, the preparation was typically viable for up to 2 hours following recovery from slicing and recordings of recurrent excitation (Fig. 2C, left trace) and inhibition (Fig. 2D right trace) could be readily obtained in voltage clamp at the reversal for inhibition and excitation, respectively.

### Viability and feasibility of motoneuron recordings from *in vitro* ventral and dorsal horn-ablated preparations

We have shown that in a semi-intact spinal cord preparation both afferent and efferent circuits can be preserved after ablation of the ventral and dorsal horn, respectively. Both types of cuts allow optical and electrophysiological access to the same dorsal pool of motoneurons, innervating the ankle flexor and extensor muscles. While we proved that recordings of ventral or dorsal root evoked synaptic currents could be obtained in both preparations, differences were observed in the conduction of the experiments across different postnatal periods: 1) with increasing age and level of myelination, cell visualization became more difficult (especially in the dorsal horn-abated preparation, in which light has to go through a thicker layer of myelin than in the case of the ventral horn-ablated cord). This increased the difficult in targeting motoneurons, but can be (at least partially) compensated if recordings are preceded by fluorescent labelling of motoneurons, either by tracing (this study) or by genetic methods. Retrograde tracing confers the further advantage of unambiguous identification of the motor nuclei innervating any muscle of choice; 2) the window of viability for recordings varies with age: preparations from older animals (regardless of the type of ablation) could be used for shorter times, typically not more than 2 hours for the mature preparations from the 4-5 weeks old weaned mice used in this study. Except for one dorsal horn-ablated preparation from a P33 mouse, we successfully obtained motoneuron recordings from all the preparations we tested at all ages (10 ventral horn and 13 dorsal horn-ablated). However, while the number of successful recordings from motoneurons was always very high in animals younger than 3 weeks (up to 17 motoneurons per cord), fewer recordings were obtained from older animals, typically between 1 and 4 per preparation. This can be attributed to the increased difficulty in visualizing and targeting cells, reduced window of viability, but also to a reduced number of healthy cells in the older preparations.

Multilevel datasets required appropriate analysis to account for data dependencies, and thus avoid erroneously treating units of analysis as independent, which may lead to a substantial rate of false positives (type I error), or the loss of statistical power and important data features when using summarized statistics (Parsons et al., 2018; Yu et al., 2022). Given the nested structure of our data (motoneuron, root, animal), we used a LMM to perform our statistical analysis, allowing us to account for, and estimate, the sources of variability. From the LMM statistical model, we will consider the estimated difference between weaned and pre-weaned groups and its 95% CI, along with an effect size (η_p_^2^), in order to infer by how much motoneuron properties or synaptic conductances might differ. The variability of each random effect is also reported inline (σ^2^ and ICC), in addition to the LMM fixed-effects estimates, to appropriately highlight the weight of each variance component in our data. Experiments obtained from dorsal horn-ablated preparations were collected with two different internal solutions (see Methods), as we later opted to use caesium based solutions for voltage clamp recordings in older animals, to improve voltage control, especially at the positive holding potentials required for recording inhibitory currents. Since the aim of this study is to establish the feasibility and utility of the described preparations, rather than providing exact quantitative measures of cell and circuit properties, we have pooled together the data obtained with different intracellular solutions (different shades of blue in Figure 3). We also noted that, with the potential exception of cell conductance, the composition of the internal solution should have little effect on the other measured parameters.

We first examined motoneuron capacitance, a reliable proxy for cell surface area, and compared recordings obtained between 2-3 weeks old (pre-weaned) and 4-5 weeks old (weaned) mice. While one would expect the cell capacitance to increase during development, in both preparations (Fig. 3A), there was no clear difference between weaned and pre-weaned mice for ventral horn (**Mean±SD**: pre-weaned: 274±78 pF, n = 61, weaned: 213±109 pF, n = 11; **LMM results**: pre-weaned_estimate_ = 272 pF, CI_95_ = [216, 328], weaned_estimated difference_ = −47pF, CI_95_ = [−142, 48]; η_p_^2^ = 0.01, CI_90_ = [0.00, 0.09]; σ^2^_Animal_ =4348 pF^2^, ICC = 0.49, σ^2^_Residuals_ = 4591 pF^2^, ICC = 0.51) and dorsal horn-ablated *in vitro* spinal cords (**Mean±SD**: pre-weaned: 261 ±84 pF, n = 69, weaned: 202±53 pF, n = 12; **LMM results**: pre-weaned_estimate_ =265 pF, CI_95_ = [224, 305], weaned_estimated difference_ = −59pF, CI_95_ = [−135, 16]; η_p_^2^ = 0.03, CI_90_ = [0.00, 0.12]; σ^2^_Animal_ = 2955 pF^2^, ICC = 0.44, σ^2^_Residuals_ = 3811 pF^2^, ICC = 0.56). This could be due to the fact that the vast majority of the recordings were performed predominantly at ages in which motoneurons’ size does not undergo any further change (Shneider et al., 2009). Despite that, we cannot exclude that some of our recordings may have been biased towards smaller motoneurons, that tend to remain healthier, especially in the more mature preparations. Regarding motoneuron conductance (Fig. 3B), despite the different intracellular solutions used, there was no difference between groups in both ventral (**Mean±SD:** pre-weaned: 43±23 nS, n = 63, weaned: 27±17, nS n = 11; **LMM results**: pre-weaned_estimate_ =41 nS, CI_95_ = [32, 51], weaned_estimated difference_ = −15 nS, CI_95_ = [−33, 4]; η_p_^2^ = 0.03, CI_90_ = [0.00, 0.13]; σ^2^_Animal_ = 104 nS^2^, ICC = 0.20, σ^2^_Residuals_ = 414 nS^2^, ICC = 0.80) and dorsal horn-ablated longitudinal preparations (**Mean±SD:** pre-weaned: 60±32 nS, n = 69, weaned: 34±11 nS, n = 10; **LMM results**: pre-weaned_estimate_ = 57 nS, CI_95_ = [43, 71], weaned_estimated difference_ = −23 nS, CI_95_ = [−51, 5]; η_p_^2^ =0.03, CI_90_ = [0.00, 0.13]; σ^2^_Animal_ = 327 nS^2^, ICC = 0.34, σ^2^_Residuals_ = 623 nS^2^, ICC = 0.66). In older mice, where the window of viability of the *in vitro* spinal cord preparations is greatly reduced, we managed to successfully obtain whole-cell recordings from healthy motoneurons. But more importantly, the range of intrinsic properties of these motoneurons indicates that they might be larger than those of previously attempted preparations from adult mice: Mitra & Brownstone (2012) report a mean motoneuron conductance of 11 nS obtained from P40-P70 mouse transverse slices, and Hadzipasic et al. (2014) report a mean motoneuron conductance up to 23nS and estimated capacitance (calculated from their reported mean motoneuron membrane time constant and mean resistance) of ~100pF, obtained from 1-6 months old mouse lumbar spinal cord slices.

Next, we compared the absolute synaptic conductance of the premotor circuits explored in this study. For sensory-related circuits, we measured monosynaptic excitation and disynaptic inhibition following dorsal root stimulation throughout the age range (Fig. 3C). Although for dorsal root-evoked excitation there was no substantial difference between weaned and preweaned mice (**Mean±SD:** pre-weaned = 26±19 nS, n = 111, pre-weaned = 15±11 nS, n = 21; **LMM results**: pre-weaned_estimate_ = 26 nS, CI_95_ = [18, 35], weaned_estimated difference_ = −11 nS, CI_95_ = [−26, 3]; η_p_^2^ = 0.02, CI_90_ = [0.00, 0.07]; σ^2^_Root_ = 34 nS^2^, ICC = 0.10, σ^2^_Animal_ = 82 nS^2^, ICC = 0.25, σ^2^_Residuals_ = 219 nS^2^, ICC = 0.65), for inhibition, the estimated absolute conductance was reduced by approximately half in 4-5 weeks old mice, with a relatively large effect size (**Mean±SD:** pre-weaned = 71±39 nS, n = 59, pre-weaned = 28±30 nS, n = 17; **LMM results:** pre-weaned_estimate_ = 72 nS, CI_95_ = [56, 88], weaned_estimated difference_ = −42 nS, CI_95_ = [−70, −14]; η_p_^2^ = 0.11, CI_90_ = [0.02, 0.23]; σ^2^_Root_ = 113 nS^2^, ICC = 0.08, σ^2^_Animal_ = 238 nS^2^, ICC = 0.17, σ^2^_Residuals_ = 1068 nS^2^, ICC = 0.75). This indicates that, while the integrity of mono-synaptic sensory pathways is fairly preserved in preparations from older animals, disynaptic afferent-related pathways might be compromised. In the dorsal horn-ablated preparation we recorded recurrent excitation and inhibition, induced by stimulation of the ventral roots, that activate the di-synaptic pathways through motoneurons and Renshaw cells respectively (Fig. 3D). Ventral root-evoked recurrent excitation was observed in all preparations throughout our age range, while in 2 animals (aged 28 and 34) we did not observe any recurrent inhibition in some root responses. In fact, while for recurrent excitation there was no striking difference between the absolute synaptic conductance of pre-weaned and weaned mice (**Mean±SD:** pre-weaned = 25±17 nS, n = 45, weaned = 13±12 nS, n = 20; **LMM results:** pre-weaned_estimate_ = 26 nS [15, 38], weaned_estimated difference_ = −12 nS, CI_95_ = [−29, 4]; η_p_^2^ = 0.03, CI_90_ = [0.00, 0.14]; σ^2^_Root_ = 15 nS^2^, ICC = 0.06, σ^2^_Animal_ = 117 nS^2^, ICC = 0.44, σ^2^_Residuals_ = 131 nS^2^, ICC = 0.50), for recurrent inhibition, this was less than 1/8 in the older group and with a large effect size (**Mean±SD:** pre-weaned = 48±33 nS, n = 124, weaned = 5±6 nS, n = 20; **LMM results:** pre-weaned_estimate_ = 48, nS, CI_95_ = [38, 57], weaned_estimated difference_ = −42 nS, CI_95_ = [−62, −22]; η_p_^2^ = 0.11, CI_90_ = [0.04, 0.20]; σ^2^_Root_ = 12 nS^2^, ICC = 0.01, σ^2^_Animal_ = 129 nS^2^, ICC = 0.14, σ^2^_Residuals_ = 819 nS^2^, ICC = 0.85). Since the recurrent inhibitory pathway is di-synaptic, its impairment could be due either to a reduced number of viable motoneurons in the preparation, thus leading to activation of a small number of Renshaw cells, or directly to a reduced number of Renshaw cells. While we cannot distinguish between these two hypotheses, it is worth pointing out that while even in older animals it was possible to record from healthy motoneurons, the impairment in a synaptic pathway that requires the recruitment of the entire motoneuron population suggests that some motoneuron loss does occur at this stage in our preparation. This impairment was larger than that of disynaptic afferent-related inhibition in 4-5 weeks old preparations, which is mediated by Ia and Ib interneurons, suggesting that the viability of pathways that do not involve motoneurons in the intermediate steps might be less affected in *in vitro* preparations from older animals. Altogether, these observations suggest that while recording cellular properties of motoneurons and their efferent and afferent inputs is feasible in 2-3 weeks old mice, the quantitative study of these pre-motor circuits after the third week of age might be at least partially impaired, especially those that require synaptic transmission through motoneurons.

## Discussion

In this work we have presented two different *in vitro* lumbar spinal cord preparations from mice, that allow electrophysiological recordings from motoneurons and preserve their afferent or efferent circuits. Both preparations can be used to obtain electrophysiological data from mice up to one month of age. Recordings became more difficult as the age increased, and anyone interested in using these preparations should carefully consider the challenges they might face when attempting *in vitro* recordings of motoneurons from ventral or dorsal horn-ablated preparations from mice older than 3 weeks. In particular, in older preparations the number of viable motoneurons might be reduced, and therefore the synaptic strengths and the degree of convergence in circuits involving motoneurons as pre-synaptic cells might be underestimated.

In the past 20 years, spinal physiologists took advantage of the fact that electrophysiological recordings from embryonic and neonatal mice *in vitro* spinal cord preparations are straightforward, thanks to tissue transparency, long viability of the preparation and resistance to anoxia and mechanical stress from slicing. In neonatal animals, whole-cell patch clamp recordings from motoneurons and spinal interneurons can be easily obtained not only from transverse slices, but also from intact or hemisected lumbar *in vitro* spinal cord preparations, where motoneurons and pre-motor circuits are fully preserved (Dyck et al., 2012; Hinckley et al., 2005; Hochman & Schmidt, 1998; Nishimaru et al., 2006). The focus on preparations from such young animals is mostly due to the technical constrains of *in vitro* preparations from more mature animals. The age range of the animals used in this study to obtain dorsal and ventral horn-ablated *in vitro* spinal cords (P15-P36), encompasses a stage when animals are already weight bearing and the locomotor pattern is established. Adult locomotor behaviours in rodents becomes most visible around 2 weeks of age, once the animal is past the weight-bearing stage and is able to fully support itself and engages into quadrupedal stepping (Altman & Sudarshan, 1975; Clarac et al., 2004). Around this stage, motoneuron collateral and sensory synapses have proliferated onto Renshaw cells and glycine becomes the sole neurotransmitter present at Renshaw-motoneuron synapses (Alvarez et al., 2013; Mentis et al., 2006; Siembab et al., 2010). Proliferation of Ia afferents onto motoneurons and other spinal interneurons such as Renshaw cells and Ia inhibitory interneurons, is also attained in 2 weeks old mice (Alvarez et al., 2013; Mentis et al., 2006, 2010, 2012; Siembab et al., 2010). Maturation of motoneuron properties also occurs in the second postnatal week (Sharples & Miles, 2021; Smith & Brownstone, 2020), which coincides with increased levels of myelination found in the lumbar cord at this stage (Foran & Peterson, 1992). Innervation patterns from descending pathways, such as corticospinal tract terminations (Donatelle, 1977) and brainstem projections (Bregman, 1987; Vinay et al., 2005) also matures by the second postnatal week. In addition, the transcriptome of proprioceptive afferents (Wu et al., 2019) and premotor spinal interneurons (Bikoff et al., 2016) remain largely unchanged from early juvenile period. Researchers interested in the study of mature spinal microcircuits will benefit from using the *in vitro* dorsal or ventral horn-ablated spinal cord from ≥2 weeks old juvenile mice. In addition to that, the accessibility to interneurons and motoneurons along with the preserved integrity of the spinal circuity, could largely benefit those interested in implementing additional techniques such as genetic manipulation and live imaging.

While we successfully managed to record from motoneurons from P15-P36 mice, cells from *in vitro* preparations obtained from ~1 month old animals were harder to target, partly because of poor optical access, and partly because the viability of the preparation was limited in time. Regarding sensory pathways, the synaptic strength of Ia excitation was not affected while disynaptic Ia/Ib inhibition was smaller in older mice (>3 weeks). On the other hand, for motor-efferent circuits, the reduced motoneuron viability in preparations from 4-5 weeks old weaned mice, may have contributed to the decreased synaptic conductances for recurrent inhibition found at this stage. This is relevant for those interested in the study of spinal circuits. We suggest the use of *in vitro* spinal cord preparations from ≥4 weeks old mice for a proof-of-concept exercise (e.g. validate a specific genetic approach), or for measuring single cell active and passive properties, but not for the quantitative comparison of synaptic strength of pre-motor circuits between a disease model and a healthy control, for instance. For this, 2-3 weeks old mice would be a more suitable choice. Another important aspect of the longitudinal *in vitro* lumbar spinal cords is that the orientation of the slicing exposes motoneurons from the dorsal nucleus of the lower lumbar cord. If researchers want to attempt these preparations from other spinal segments (e.g. thoracic, cervical) of late postnatal mice, they must carefully take into consideration the anatomy of the motor nuclei for efficient coronal slicing. In other areas of the nervous system, such as the brainstem, different *in vitro* electrophysiological preparations have been developed by using anatomical cues and different angles of slicing, in order to gain access to specific nuclei (Anderson et al., 2016; Ruangkittisakul et al., 2014). Therefore, one may adopt and morph some of the methodology used in this study, in order to develop novel *in vitro* preparations that might grant access to other pathways and motor nuclei within the mouse spinal cord.

The data generated from our *in vitro* recordings have a hierarchical structure, which is a common feature in neuroscience datasets (Aarts et al., 2014). In our experimental setup, motoneuron were recorded from different animals (ranging from 1 to 17 cells per preparation) and synaptic responses were obtained following L4 and L5 dorsal or ventral root stimulations from different *in vitro* spinal cords (up to 4 different roots per animal). Lately, a greater focus has been put on the use of inappropriate statistical methods in basic neuroscience research that do not take into consideration data dependencies, resulting in a higher rate of type I error (false positives) (Aarts et al., 2014; Parsons et al., 2018; Yu et al., 2022). This issue has been recently raised regarding data obtained from multiple *in vivo* motoneuron recordings obtained from different animals (Baczyk, 2022; Baczyk et al., 2020; Héroux, 2021). Different statistical approaches can be used to account for data dependencies such as the hierarchical bootstraps and LMMs. The hierarchical bootstrap performs ideally with large samples sizes, such as those containing hundreds of observations per each animal, and therefore would not be the most appropriate for the datasets we generated since the resampling procedure would not accurately represent the population distribution of motoneuron intrinsic properties or synaptic conductances we intend to study (Saravanan et al., 2020). LMMs are a suitable approach to nested designs since they allow to estimate and account for variance across different levels, and have a small false positive rate even for small sample sizes (Saravanan et al., 2020; Yu et al., 2022). We therefore decided to use a LMM to compare intrinsic motoneuron properties and synaptic conductances obtained from pre-weaned and weaned mice. The model provides an estimate and respective CI for the control group (pre-weaned) and the difference between groups (weaned minus pre-weaned) along with the variance contained within each level of the data structure (e.g. animal, root, synaptic response). In addition we also calculated an effect size (η_p_^2^), that provides a better understanding of the magnitude of any biological differences between groups (Cohen, 1988; Richardson, 2011). We based a large portion of the discussion of this innovative methodology paper on some of the variables from our statistical analysis (summarized in Supplemental Table 1), which provided a deeper understanding of the viability of motoneuron recordings and integrity of pre-motor circuits across age. Regarding the variance component of our hierarchical datasets, we found that, in some instances, motoneuron intrinsic properties and synaptic responses to root stimulation contributed to ~50% of the total variability found in our datasets (measured as ICC from σ^2^_Residuals_), which likely reflects the highly heterogeneous properties of motoneurons (Highlander et al., 2020). However this was not the case for all parameters, as some showed high variability within the lowest level of the LMM structure (σ^2^_Residuals_), ranging from 65% to 85% of total variance (see Supplemental Table 1). This is relevant for experimental nested designs involving recordings from multiple motoneurons from the same animal and repeating the experiments on different animals. This design inevitably gives rise to data with a nested structure, and therefore appropriate models that take into consideration dependence in multilevel data structures, such as LMM, should be routinely employed to understand and take into account the sources of variability in these type of datasets (Yu et al., 2022).

Since the first use of an isolated spinal cord preparation almost 50 years ago (Otsuka & Konihsi, 1974), spinal physiologists have been largely constrained to the use of *in vitro* preparations from neonatal mice to interrogate the cellular mechanisms of the locomotor control systems within the mammalian spinal cord. The longitudinal spinal cords from this study are therefore a suitable tool to those interested in studying motor circuits, since viable preparations can be easily obtained from 2-3 weeks old animals, whose locomotion is similar or indistinguishable from that of adults, and that have reached full circuit assembly and maturation. The thickness of the *in vitro* preparation will also allow a much better study of circuits and motoneuron properties without significant adverse effects from slicing (Mousa & Elbasiouny, 2021). The *in vitro* preparations described in this work should therefore captivate the interest of those interested in the study of mammalian spinal networks, since they provide a much needed and useful choice to probe motoneurons and their pre-motor circuits.

## Data Availability

Data are made available in the supporting files.

## Author Contributions

MGO, MB and FN conceived and designed research

MGO, JOA, MB and FN performed experiments

MGO and FN analysed data

MGO, MB and FN interpreted results of experiments

MGO, MB and FN prepared figures

MGO, MB and FN drafted manuscript

MGO, JOA, MB and FN edited and revised manuscript

MGO, JOA, MB and FN approved final version of manuscript.

---

**Supplementary figure 1.**
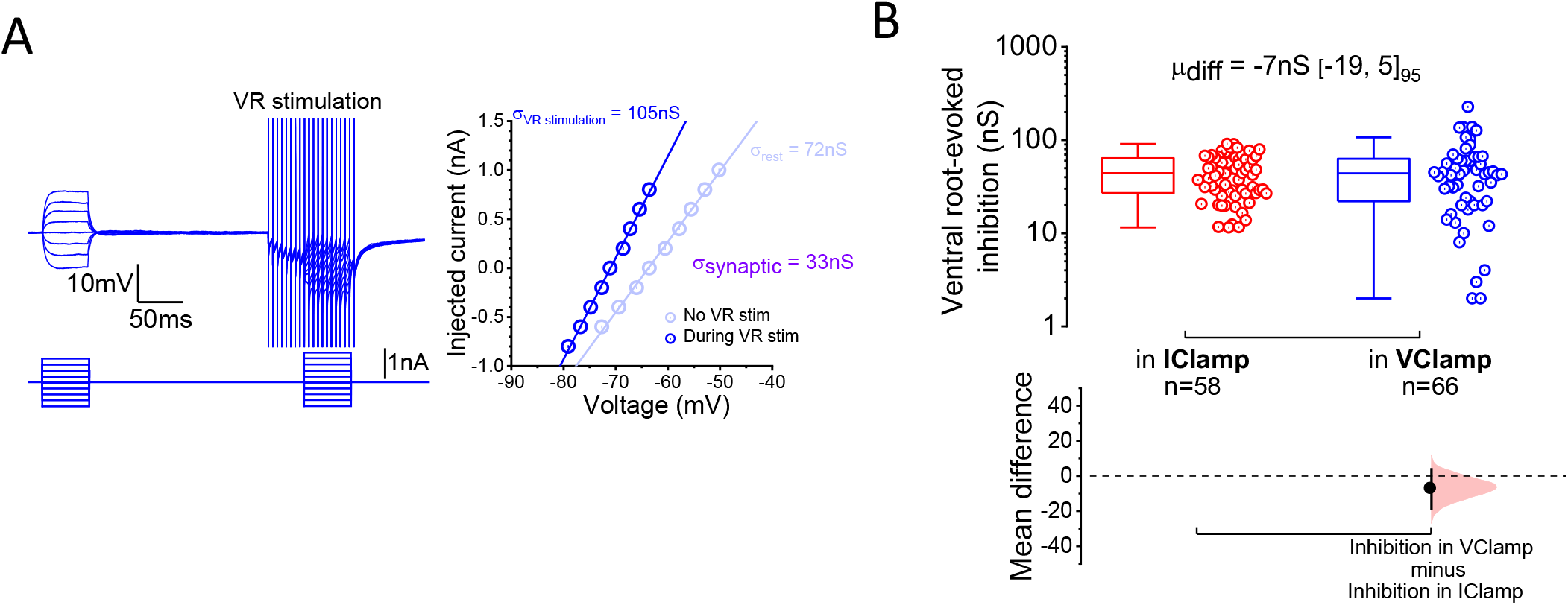
Recurrent inhibition estimated in current clamp. **(a)** Example of a voltage response from a motoneuron to current injection before and during high-frequency (200Hz for 100ms) ventral root stimulation (left), with respective current-voltage plot used to estimate synaptic conductance (right). **(b)** Estimation plot comparing the synaptic conductance for recurrent inhibition estimated in current (IClamp) and voltage clamp (VClamp). Each “n” is a L4 or L5 ventral root-evoked synaptic response obtained from a motoneuron. The bootstrapped mean difference (μ_diff_) and 95% CI are shown at the top of the plot.

**Supplemental Table 1.**
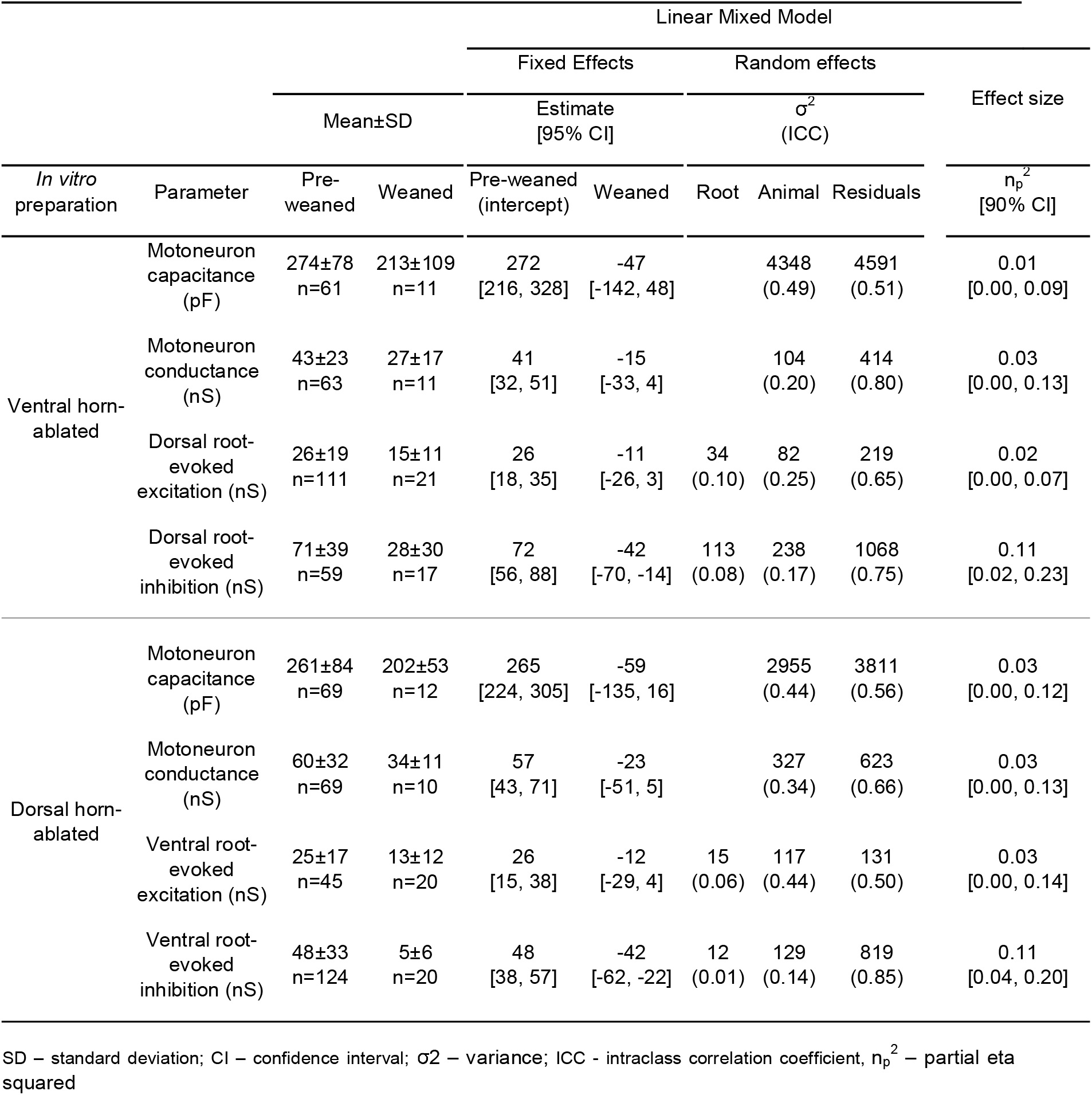
Mean and standard deviation, linear mixed model fixed and random-effects variables and partial eta squared for motoneuron capacitance, motoneuron conductance and synaptic conductances from ventral and dorsal horn-ablated *in vitro* spinal cord preparations

## References

Aarts, E., Verhage, M., Veenvliet, J. V, Dolan, C. V, & Sluis, S. Van Der. (2014). A solution to dependency□: using multilevel analysis to accommodate nested data. 17(4), 491–496. https://doi.org/10.1038/nn.3648

Altman, J., & Sudarshan, K. (1975). Postnatal development of locomotion in the laboratory rat. Animal Behaviour, 23(PART 4), 896–920. https://doi.org/10.1016/0003-3472(75)90114-1

Alvarez, F. J., Benito-Gonzalez, A., & Siembab, V. C. (2013). Principles of interneuron development learned from Renshaw cells and the motoneuron recurrent inhibitory circuit. Annals of the New York Academy of Sciences, 1279(1), 22–31. https://doi.org/10.1111/nyas.12084

Anderson, T. M., Garcia, A. J., Baertsch, N. A., Pollak, J., Bloom, J. C., Wei, A. D., Rai, K. G., & Ramirez, J. M. (2016). A novel excitatory network for the control of breathing. Nature, 536(7614), 76–80. https://doi.org/10.1038/nature18944

Arber, S. (2012). Motor Circuits in Action: Specification, Connectivity, and Function. Neuron, 74(6), 975–989. https://doi.org/10.1016/j.neuron.2012.05.011

Baczyk, M. (2022). Reply to Dr. Heroux. Journal of Applied Physiology, December 2020, 8750. https://doi.org/10.1113/JP280274.Alami

Baczyk, M., Drzymała-Celichowska, H., Mrówczyński, W., & Krutki, P. (2020). Polarity-dependent adaptations of motoneuron electrophysiological properties after 5-wk transcutaneous spinal direct current stimulation in rats. Journal of Applied Physiology, 129(4), 646–655. https://doi.org/10.1152/japplphysiol.00301.2020

Bagust, J., Forsythe, I. d., Kerkut, G. A., & Loots, J. M. (1982). Synaptic and non-synaptic components of the dorsal horn potential in isolated hamster spinal cord. Brain Research, 233(1), 186–194. https://doi.org/10.1016/0006-8993(82)90940-4

Bagust, J., & Kerkut, G. A. (1979). Some effects of magnesium ions upon conduction and synaptic activity in the isolated spinal cord of the mouse. Brain Research, 177(2), 410–413. https://doi.org/10.1016/0006-8993(79)90796-0

Bates, D., Mächler, M., Bolker, B. M., & Walker, S. C. (2015). Fitting linear mixed-effects models using lme4. Journal of Statistical Software, 67(1). https://doi.org/10.18637/jss.v067.i01

Bhumbra, G. S., & Beato, M. (2018). Recurrent excitation between motoneurones propagates across segments and is purely glutamatergic. PLoS Biology, 16(3), 1–16. https://doi.org/10.1371/journal.pbio.2003586

Bikoff, J. B., Gabitto, M. I., Rivard, A. F., Drobac, E., MacHado, T. A., Miri, A., Brenner-Morton, S., Famojure, E., Diaz, C., Alvarez, F. J., Mentis, G. Z., & Jessell, T. M. (2016). Spinal Inhibitory Interneuron Diversity Delineates Variant Motor Microcircuits. Cell, 165(1), 207–219. https://doi.org/10.1016/j.cell.2016.01.027

Bregman, B. S. (1987). Development of serotonin immunoreactivity in the rat spinal cord and its plasticity after neonatal spinal cord lesions. Developmental Brain Research, 34(2), 245–263. https://doi.org/10.1016/0165-3806(87)90213-6

Brink, B. Y. E., Jankowska, E., Mccreat, D. A., & Skoog, B. (1983). Inhibitory interactions between interneurones in reflex pathways from group Ia and group Ib afferents in the cat. Journal of Physiology, 361–373.

Brownstone, R. M. (2006). Beginning at the end: Repetitive firing properties in the final common pathway. Progress in Neurobiology, 78(3–5), 156–172. https://doi.org/10.1016/j.pneurobio.2006.04.002

Clarac, F., Brocard, F., & Vinay, L. (2004). The maturation of locomotor networks. Progress in Brain Research, 143(03), 57–66. https://doi.org/10.1016/S0079-6123(03)43006-9

Coe, R. (2002). Effect Size guide. 1–18.

Cohen, J. (1988). Statistical Power Analysis for the Behavioral Sciences. Routledge.

Correll, J., Mellinger, C., & Pedersen, E. J. (2021). Flexible approaches for estimating partial eta squared in mixed-effects models with crossed random factors. Behavior Research Methods. https://doi.org/10.3758/s13428-021-01687-2

Donatelle, J. M. (1977). Growth of the corticospinal tract and the development of placing reactions in the postnatal rat. Journal of Comparative Neurology, 175(2), 207–231. https://doi.org/10.1002/cne.901750205

Dougherty, K. J., & Kiehn, O. (2010). Firing and cellular properties of V2a interneurons in the rodent spinal cord. Journal of Neuroscience, 30(1), 24–37. https://doi.org/10.1523/JNEUROSCI.4821-09.2010

Dugué, G. P., Dumoulin, A., Triller, A., & Dieudonné, S. (2005). Target-dependent use of coreleased inhibitory transmitters at central synapses. Journal of Neuroscience, 25(28), 6490–6498. https://doi.org/10.1523/JNEUROSCI.1500-05.2005

Dyck, J., Lanuza, G. M., & Gosgnach, S. (2012). Functional characterization of dI6 interneurons in the neonatal mouse spinal cord. Journal of Neurophysiology, 107(12), 3256–3266. https://doi.org/10.1152/jn.01132.2011

Eccles, J. C. (1957). The Physiology of Nerve Cells. John Hopkins Press.

Foran, D. R., & Peterson, A. C. (1992). Myelin acquisition in the central nervous system of the mouse revealed by an MBP-Lac Z transgene. Journal of Neuroscience, 12(12), 4890–4897. https://doi.org/10.1523/jneurosci.12-12-04890.1992

Goulding, M. (2009). Circuits controlling vertebrate locomotion: moving in a new direction. Nat Rev Neurosci, 10(7), 507–518. https://doi.org/10.1038/nrn2608

Hadzipasic, M., Tahvildari, B., Nagy, M., Bian, M., Horwich, A. L., & McCormick, D. A. (2014). Selective degeneration of a physiological subtype of spinal motor neuron in mice with SOD1-linked ALS. Proceedings of the National Academy of Sciences of the United States of America, 111(47), 16883–16888. https://doi.org/10.1073/pnas.1419497111

Hamill, P. O., Marty, A., Neher, A., B., S., & Sigworth, F. J. (1981). Improved patch-clamp techniques for high-resolution current recording from cells and cell-free membrane patches. Pflugers Arch., 391(2), 85–100. https://doi.org/10.1007/BF00656997

Henderson, C. R., Kempthorne, O., Searle, S. R., & Krosigk, C. M. (1959). The Estimation of Environmental and Genetic Trends from Records Subject to Culling. Biometrics, 15(2), 192–218.

Héroux, M. E. (2021). Analyzing dependent data as if independent biases effect size estimates and increases the risk of false-positive findings. Journal of Applied Physiology, 130(3), 675–676. https://doi.org/10.1152/JAPPLPHYSIOL.01024.2020

Highlander, M. M., Allen, J. M., & Elbasiouny, S. M. (2020). Meta-analysis of biological variables’ impact on spinal motoneuron electrophysiology data. Journal of Neurophysiology, 123(4), 1380–1391. https://doi.org/10.1152/jn.00378.2019

Hinckley, C. A., Hartley, R., Wu, L., Todd, A., & Ziskind-Conhaim, L. (2005). Locomotor-like rhythms in a genetically distinct cluster of interneurons in the mammalian spinal cord. Journal of Neurophysiology, 93(3), 1439–1449. https://doi.org/10.1152/jn.00647.2004

Ho, J., Tumkaya, T., Aryal, S., Choi, H., & Claridge-Chang, A. (2019). Moving beyond P values: data analysis with estimation graphics. Nature Methods, 16(7), 565–566. https://doi.org/10.1038/s41592-019-0470-3

Hochman, S., & Schmidt, B. J. (1998). Whole cell recordings of lumbar motoneurons during locomotor-like activity in the in vitro neonatal rat spinal cord. Journal of Neurophysiology, 79(2), 743–752. https://doi.org/10.1152/jn.1998.79.2.743

Hultborn, H. (2006). Spinal reflexes, mechanisms and concepts: From Eccles to Lundberg and beyond. Progress in Neurobiology, 78(3–5), 215–232. https://doi.org/10.1016/j.pneurobio.2006.04.001

Hultborn, H., & Udo, M. (1972). Convergence on interneurones in the reciprocal Ia inhibitory pathway to motoneurones. Acta Physiologica Scandinavica, 85, 1–42. https://doi.org/10.1111/j.1748-1716.1972.tb05298.x

Jensen, D. B., Kadlecova, M., Allodi, I., & Meehan, C. F. (2021). Response to Letter to Editor on the article Jensen DB, Kadlecova M, Allodi I, Meehan CF (2020). Journal of Physiology, 599(17), 4233–4236. https://doi.org/10.1113/JP281539

Jessell, T. M. (2000). Neuronal specification in the spinal cord: inductive signals and transcriptional codes. Nature, 20(1), 20–29.

Li, E. Z., Garcia-Ramirez, D. L., & Dougherty, K. J. (2019). Flexor and Extensor Ankle Afferents Broadly Innervate Locomotor Spinal Shox2 Neurons and Induce Similar Effects in Neonatal Mice. Frontiers in Cellular Neuroscience, 13(October), 1–18. https://doi.org/10.3389/fncel.2019.00452

Manuel, M. (2021). Suboptimal discontinuous current-clamp switching rates lead to deceptive mouse neuronal firing. ENeuro, 8(1), 1–14. https://doi.org/10.1523/ENEURO.0461-20.2020

Manuel, M., Iglesias, C., Donnet, M., Leroy, F., Heckman, C. J., & Zytnicki, D. (2009). Fast Kinetics, High-Frequency Oscillations, and Subprimary Firing Range in Adult Mouse Spinal Motoneurons. The Journal of Neuroscience, 29(36), 11246–11256. https://doi.org/10.1523/JNEUROSCI.3260-09.2009

Manuel, M., & Zytnicki, D. (2021). Comments on the article by Jensen et al. (2020). Journal of Physiology, 599(17), 4231–4232. https://doi.org/10.1113/JP281461

Meehan, C. F., Sukiasyan, N., Zhang, M., Nielsen, J. B., & Hultborn, H. (2010). Intrinsic properties of mouse lumbar motoneurons revealed by intracellular recording in vivo. Journal of Neurophysiology, 103(5), 2599–2610. https://doi.org/10.1152/jn.00668.2009

Meehan, Claire F., Mayr, K. A., Manuel, M., Nakanishi, S. T., & Whelan, P. J. (2017). Decerebrate mouse model for studies of the spinal cord circuits. Nature Protocols, 12(4), 732–747. https://doi.org/10.1038/nprot.2017.001

Mentis, G. Z., Alvarez, F. J., Shneider, N. A., Siembab, V. C., & O’Donovan, M. J. (2010). Mechanisms regulating the specificity and strength of muscle afferent inputs in the spinal cord. Annals of the New York Academy of Sciences, 1198, 220–230. https://doi.org/10.1111/j.1749-6632.2010.05538.x

Mentis, G. Z., Blivis, D., Liu, W., Drobac, E., Melissa, E., Kong, L., Alvarez, F. J., Sumner, C. J., Michael, J., & Donovan, O. (2012). Early functional impairment of sensory-motor connectivity in a mouse model of spinal muscular atrophy. Neuron, 69(3), 453–467. https://doi.org/10.1016/j.neuron.2010.12.032.Early

Mentis, G. Z., Siembab, V. C., Zerda, R., O’Donovan, M. J., & Alvarez, F. J. (2006). Primary afferent synapses on developing and adult Renshaw cells. Journal of Neuroscience, 26(51), 13297–13310. https://doi.org/10.1523/JNEUROSCI.2945-06.2006

Mitra, P., & Brownstone, R. M. (2012). An in vitro spinal cord slice preparation for recording from lumbar motoneurons of the adult mouse. Journal of Neurophysiology, 107(2), 728–741. https://doi.org/10.1152/jn.00558.2011

Mousa, M. H., & Elbasiouny, S. M. (2021). Estimating the effects of slicing on the electrophysiological properties of spinal motoneurons under normal and disease conditions. Journal of Neurophysiology, 125(4), 1450–1467. https://doi.org/10.1152/jn.00543.2020

Nishimaru, H., Restrepo, C. E., & Kiehn, O. (2006). Activity of Renshaw cells during locomotor-like rhythmic activity in the isolated spinal cord of neonatal mice. Journal of Neuroscience, 26(20), 5320–5328. https://doi.org/10.1523/JNEUROSCI.5127-05.2006

O’Donovan, M., Ho, S., & Yee, W. (1994). Calcium imaging of rhythmic network activity in the developing spinal cord of the chick embryo. Journal of Neuroscience, 14(11 I), 6354–6369. https://doi.org/10.1523/jneurosci.14-11-06354.1994

Otsuka, M., & Konihsi, S. (1974). Electrophysiology of mammalian spinal cord in vitro. Nature, 252, 733–734.

Parsons, N. R., Teare, M. D., & Sitch, A. J. (2018). Unit of analysis issues in laboratory-based research. ELife, 7, 1–25. https://doi.org/10.7554/eLife.32486

Renshaw, B. (1946). Central effects of centripetal impulses in axons of spinal ventral roots. Journal of Neurophysiology, 9, 191–204.

Richardson, J. T. E. (2011). Eta squared and partial eta squared as measures of effect size in educational research. Educational Research Review, 6(2), 135–147. https://doi.org/10.1016/j.edurev.2010.12.001

Ruangkittisakul, A., Kottick, A., Picardo, M. C. D., Ballanyi, K., & Del Negro, C. A. (2014). Identification of the pre-Bötzinger complex inspiratory center in calibrated “sandwich” slices from newborn mice with fluorescent Dbx1 interneurons. Physiological Reports, 2(8), 1–16. https://doi.org/10.14814/phy2.12111

Saravanan, V., Berman, G. J., & Sober, S. J. (2020). Application of the hierarchical bootstrap to multi-level data in neuroscience. Neurons, Behavior, Data Analysis and Theory, 3(5), 1–29. http://www.ncbi.nlm.nih.gov/pubmed/33644783%0Ahttp://www.pubmedcentral.nih.gov/articlerender.fcgi?artid=PMC7906290

Sharples, S. A., & Miles, G. B. (2021). Maturation of persistent and hyperpolarization-activated inward currents shape the recruitment of motoneuron subtypes during postnatal development. ELife. https://doi.org/10.7554/eLife.71385

Sherrington, C. S. (1906). The integrative action of the nervous system. Yale University Press.

Shneider, N. A., Brown, M. N., Smith, C. A., Pickel, J., & Alvarez, F. J. (2009). Gamma motor neurons express distinct genetic markers at birth and require muscle spindle-derived GDNF for postnatal survival. Neural Development, 4(1). https://doi.org/10.1186/1749-8104-4-42

Siembab, V. C., Smith, C. A., Zagoraiou, L., Berrocal, M. C., Mentis, G. Z., & Alvarez, F. J. (2010). Target selection of proprioceptive and motor axon synapses on neonatal V1-derived Ia inhibitory interneurons and Renshaw cells. Journal of Comparative Neurology, 518(23), 4675–4701. https://doi.org/10.1002/cne.22441

Smith, C. C., & Brownstone, R. M. (2020). Spinal motoneuron firing properties mature from rostral to caudal during postnatal development of the mouse. Journal of Physiology, 598(23), 5467–5485. https://doi.org/10.1113/JP280274

Steiger, J. H. (2004). Beyond the F test: Effect size confidence intervals and tests of close fit in the analysis of variance and contrast analysis. Psychological Methods, 9(2), 164–182. https://doi.org/10.1037/1082-989X.9.2.164

Takahashi, T. (1990). Membrane currents in visually identified motoneurones of neonatal rat spinal cord. The Journal of Physiology, 423(1), 27–46. https://doi.org/10.1113/jphysiol.1990.sp018009

Vinay, L., Ben-Mabrouk, F., Brocard, F., Clarac, F., Jean-Xavier, C., Pearlstein, E., & Pflieger, J. F. (2005). Perinatal development of the motor systems involved in postural control. Neural Plasticity, 12(2–3), 131–139. https://doi.org/10.1155/NP.2005.131

Wang, Z., Li, L., Goulding, M., Frank, E., Wang, Z., Ly, L., Goulding, M., & Early, F. E. (2021). Early Postnatal Development of Reciprocal Ia Inhibition in the Murine Spinal Cord. 185–196. https://doi.org/10.1152/jn.90354.2008.

Whelan, P. J. (2003). Developmental aspects of spinal locomotor function: Insights from using the in vitro mouse spinal cord preparation. Journal of Physiology, 553(3), 695–706. https://doi.org/10.1113/jphysiol.2003.046219

Wilson, J. M., Dombeck, D. A., Díaz-Ríos, M., Harris-Warrick, R. M., & Brownstone, R. M. (2007). Two-photon calcium imaging of network activity in XFP-expressing neurons in the mouse. Journal of Neurophysiology, 97(4), 3118–3125. https://doi.org/10.1152/jn.01207.2006

Wu, D., Schieren, I., Qian, Y., Zhang, C., Jessell, T. M., & De Nooij, J. C. (2019). A role for sensory end organ-derived signals in regulating muscle spindle proprioceptor phenotype. Journal of Neuroscience, 39(22), 4252–4267. https://doi.org/10.1523/JNEUROSCI.2671-18.2019

Yu, Z., Guindani, M., Grieco, S. F., Chen, L., Holmes, T. C., & Xu, X. (2022). Beyond t test and ANOVA: applications of mixed-effects models for more rigorous statistical analysis in neuroscience research. Neuron, 110(1), 21–35. https://doi.org/10.1016/j.neuron.2021.10.030

